# Bioconda: A sustainable and comprehensive software distribution for the life sciences

**DOI:** 10.1101/207092

**Authors:** Björn Grüning, Ryan Dale, Andreas Sjödin, Brad A. Chapman, Jillian Rowe, Christopher H. Tomkins-Tinch, Renan Valieris, Adam Caprez, Bérénice Batut, Mathias Haudgaard, Thomas Cokelaer, Kyle A. Beauchamp, Brent S Pedersen, Youri Hoogstrate, Anthony Bretaudeau, Devon Ryan, Gildas Le Corguillé, Dilmurat Yusuf, Sebastian Luna-Valero, Rory Kirchner, Karel Brinda, Thomas Wollmann, Martin Raden, Simon J. van Heeringen, Nicola Soranzo, Lorena Pantano, Zachary Charlop-Powers, Per Unneberg, Matthias De Smet, Marcel Martin, Greg Von Kuster, Tiago Antao, Milad Miladi, Kevin Thornton, Christian Brueffer, Marius van den Beek, Daniel Maticzka, Clemens Blank, Sebastian Will, K´evin Gravouil, Joachim Wolff, Manuel Holtgrewe, Jörg Fallmann, Vitor C. Piro, Ilya Shlyakhter, Ayman Yousif, Philip Mabon, Xiao-Ou Zhang, Wei Shen, Jennifer Cabral, Cristel Thomas, Eric Enns, Joseph Brown, Jorrit Boekel, Mattias de Hollander, Jerome Kelleher, Nitesh Turaga, Julian R. de Ruiter, Dave Bouvier, Simon Gladman, Saket Choudhary, Nicholas Harding, Florian Eggenhofer, Arne Kratz, Zhuoqing Fang, Robert Kleinkauf, Henning Timm, Peter J. A. Cock, Enrico Seiler, Colin Brislawn, Hai Nguyen, Endre Bakken Stovner, Philip Ewels, Matt Chambers, James E. Johnson, Emil Hägglund, Simon Ye, Roman Valls Guimera, Elmar Pruesse, W. Augustine Dunn, Lance Parsons, Rob Patro, David Koppstein, Elena Grassi, Inken Wohlers, Alex Reynolds, MacIntosh Cornwell, Nicholas Stoler, Daniel Blankenberg, Guowei He, Marcel Bargull, Alexander Junge, Rick Farouni, Mallory Freeberg, Sourav Singh, Daniel R. Bogema, Fabio Cumbo, Liang-Bo Wang, David E Larson, Matthew L. Workentine, Upendra Kumar Devisetty, Sacha Laurent, Pierrick Roger, Xavier Garnier, Rasmus Agren, Aziz Khan, John M Eppley, Wei Li, Bianca Katharina Stöcker, Tobias Rausch, James Taylor, Patrick R. Wright, Adam P. Taranto, Davide Chicco, Bengt Sennblad, Jasmijn A. Baaijens, Matthew Gopez, Nezar Abdennur, Iain Milne, Jens Preussner, Luca Pinello, Avi Srivastava, Aroon T. Chande, Philip Reiner Kensche, Yuri Pirola, Michael Knudsen, Ino de Bruijn, Kai Blin, Giorgio Gonnella, Oana M. Enache, Vivek Rai, Nicholas R. Waters, Saskia Hiltemann, Matthew L. Bendall, Christoph Stahl, Alistair Miles, Yannick Boursin, Yasset Perez-Riverol, Sebastian Schmeier, Erik Clarke, Kevin Arvai, Matthieu Jung, Tom´as Di Domenico, Julien Seiler, Eric Rasche, Etienne Kornobis, Daniela Beisser, Sven Rahmann, Alexander S Mikheyev, Camy Tran, Jordi Capellades, Christopher Schröder, Adrian Emanuel Salatino, Simon Dirmeier, Timothy H. Webster, Oleksandr Moskalenko, Gordon Stephen, Johannes Köster

## Abstract

We present Bioconda (https://bioconda.github.io), a distribution of bioinformatics software for the lightweight, multiplatform and language-agnostic package manager Conda. Currently, Bioconda offers a collection of over 3000 software packages, which is continuously maintained, updated, and extended by a growing global community of more than 200 contributors. Bioconda improves analysis reproducibility by allowing users to define isolated environments with defined software versions, all of which are easily installed and managed without administrative privileges.

## Introduction

Thousands of new software tools have been released for bioinformatics in recent years, in a variety of programming languages. Accompanying this diversity of construction is an array of installation methods. Often, Software has to be compiled manually for different hardware architectures and operating systems, with management left to the user or system administrator. Scripting languages usually deliver their own package management tools for installing, updating, and removing packages, though these are often limited in scope to packages written in the same scripting language and cannot handle external dependencies (e.g., C libraries). Published scientific software often consists of simple collections of custom scripts distributed with textual descriptions of the manual steps required to install the software. New analyses often require novel combinations of multiple tools, and the heterogeneity of scientific software makes management of a software stack complicated and error-prone. Moreover, it inhibits reproducible science (Mesirov, 2010; Baker, 2016; Munafò et al., 2017), because it is hard to reproduce a software stack on different machines. System-wide deployment of software has traditionally been handled by administrators, but reproducibility often requires that the researcher (who is often not an expert in administration) is able to maintain full control of the software environment and rapidly modify it without administrative privileges.

The Conda package manager (https://conda.io) has become an increasingly popular approach to overcome these challenges. Conda normalizes software installations across language ecosystems by describing each software package with a *recipe* that defines meta-information and dependencies, as well as a *build script* that performs the steps necessary to build and install the software. Conda prepares and builds software packages within an isolated environment, transforming them into relocatable binaries. Conda packages can be built for all three major operating systems: Linux, macOS, and Windows. Importantly, installation and management of packages requires no administrative privileges, such that a researcher can control the available software tools regardless of the underlying infrastructure. Moreover, Conda obviates reliance on system-wide installation by allowing users to generate isolated software environments, within which versions and tools can be managed per-project, without generating conflicts or incompatibilities (see online methods). These environments support reproducibility, as they can can be rapidly exchanged via files that describe their installation state. Conda is tightly integrated into popular solutions for reproducible scientific data analysis like Galaxy (Afgan et al., 2016), bcbio-nextgen (https://github.com/chapmanb/bcbio-nextgen), and Snakemake (Köster and Rahmann, 2012). Finally, while Conda provides many commonly-used packages by default, it also allows users to optionally include additional repositories (termed *channels)* of packages that can be installed.

## Results

In order to unlock the benefits of Conda for the life sciences, the Bioconda project was founded in 2015. The mission of Bioconda is to make bioinformatics software easily installable and manageable via the Conda package manager. Via its channel for the Conda package manager, Bioconda currently provides over 3000 software packages for Linux and macOS. Development is driven by an open community of over 200 international scientists. In the prior two years, package count and the number of contributors have increased linearly, on average, with no sign of saturation (Fig. 1a,b). The barrier to entry is low, requiring a willingness to participate and adherence to community guidelines. Many software developers contribute recipes for their own tools, and many Bioconda contributors are invested in the project as they are also users of Conda and Bioconda. Bioconda provides packages from various language ecosystems like Python, R (CRAN and Bioconductor), Perl, Haskell, as well as a plethora of C/C++ programs (Fig. 1c). Many of these packages have complex dependency structures that require various manual steps to install when not relying on a package manager like Conda (Fig. 2a, Online Methods). With over 6.3 million downloads, the service has become a backbone of bioinformatics infrastructure (Fig. 1d). Bioconda is complemented by the conda-forge project (https://conda-forge.github.io), which hosts software not specifically related to the biological sciences. The two projects collaborate closely, and the Bioconda team maintains over 500 packages hosted by conda-forge. Among all currently available distributions of bioinformatics software, Bioconda is by far the most comprehensive, while being among the youngest (Fig. 2d).

**Figure 1:**
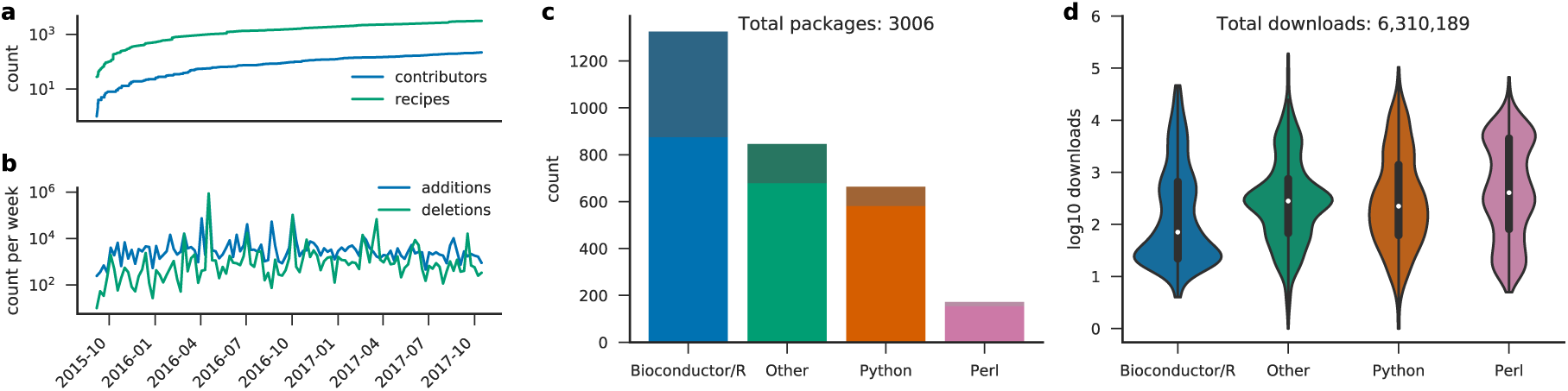
Bioconda development and usage since the beginning of the project. (a) contributing authors and added recipes over time. (b) code line additions and deletions per week. (c) package count per language ecosystem (saturated colors on bottom represent explicitly life science related packages). (d) total downloads per language ecosystem. The term “other” entails all recipes that do not fall into one of the specific categories. Note that a subset of packages that started in Bioconda have since been migrated to the more appropriate, general-purpose conda-forge channel. Older versions of such packages still reside in the Bioconda channel, and as such are included in the recipe count (a) and download count (d). Statistics obtained Oct. 25, 2017.

**Figure 2:**
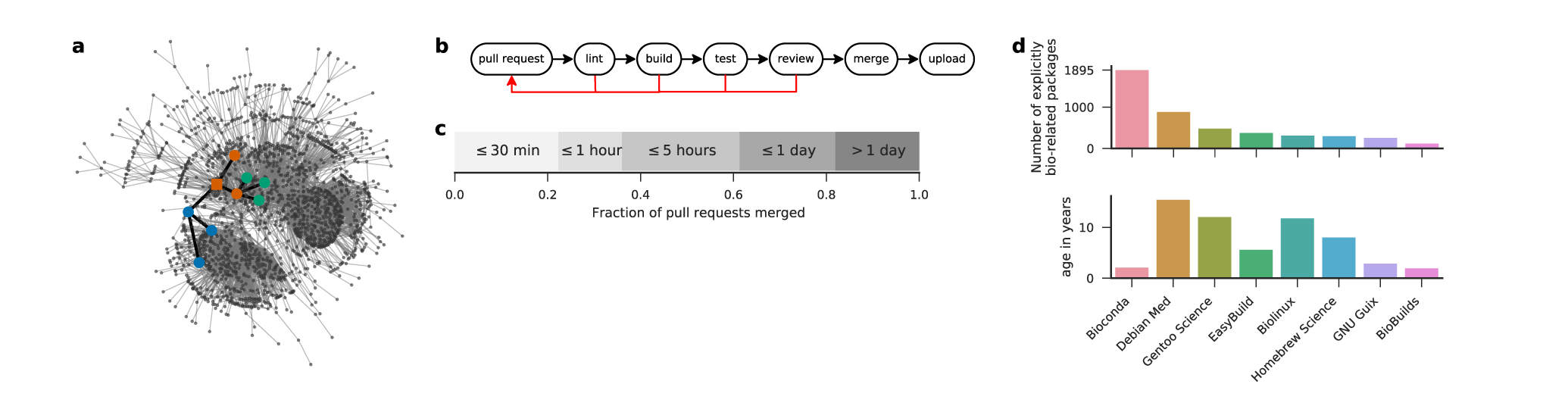
Dependency structure, workflow, comparison with other resources, and turnaround time. (a) largest connected component of directed acyclic graph of Bioconda packages (nodes) and dependen- cies (edges). Highlighted is the induced subgraph of the CNVkit (Talevich et al., 2016) package and it’s dependencies (node coloring as defined in Fig. 1c, squared node represents CNVkit). (b) GitHub based development workflow: a contributor provides a pull request that undergoes several build and test steps, followed by a human review. If any of these checks does not succeed, the contributor can update the pull request accordingly. Once all steps have passed, the changes can be merged. (c) Turnaround time from submission to merge of pull requests in Bioconda. (d) Comparison of explicitly life science related packages in Bioconda with Debian Med (https://www.debian.org/devel/debian-med), Gentoo Science Overlay (category sci-biology, https://github.com/gentoo/sci), EasyBuild (module bio, https://easybuilders.github.io/easybuild), Biolinux (Field et al., 2006), Homebrew Science (tag bioinformatics, https://brew.sh), GNU Guix (category bioinformatics, https://www.gnu.org/s/guix), and BioBuilds (https://biobuilds.org). The lower panel shows the project age since the first release or commit. Statistics obtained Oct. 23, 2017.

To ensure reliable maintenance of such numbers of packages, we use a semi-automatic, agent-assisted development workflow (Fig. 2b). All Bioconda recipes are hosted in a GitHub repository (https://github.com/bioconda/bioconda-recipes). Both the addition of new recipes and the update of existing recipes in Bioconda is handled via *pull requests*. Thereby, a modified version of one or more recipes is compared against the current state of Bioconda. Once a pull request arrives, our infrastructure performs several automatic checks. Problems discovered in any step are reported to the contributor and further progress is blocked until they are resolved. First, the modified recipes are checked for syntactic anti-patterns, i.e., formulations that are syntactically correct but bad style (termed *linting*). Second, the modified recipes are built on Linux and macOS, via a cloud based, free-of-charge service (https://travis-ci.org). Successfully built recipes are tested (e.g., by running the generated executable). Since Bioconda packages must be able to run on any supported system, it is important to check that the built packages do not rely on particular elements from the build environment. Therefore, testing happens in two stages: (a) test cases are executed in the build environment (b) test cases are executed in a minimal Docker (https://docker.com) container which purposefully lacks all non-common system libraries (hence, a dependency that is not explicitly defined will lead to a failure). Once the *build* and *test* steps have succeeded, a member of the Bioconda team reviews the proposed changes and, if acceptable, merges the modifications into the official repository. Upon merging, the recipes are built again and uploaded to the hosted Bioconda channel (https://anaconda.org/bioconda), where they become available via the Conda package manager. When a Bioconda package is updated to a new version, older builds are generally preserved, and recipes for multiple older versions may be maintained in the Bioconda repository. The usual turnaround time of above workflow is short (Fig. 2d). 61% of the pull requests are merged within 5 hours. Of those, 36% are even merged within 1 hour. Only 18% of the pull requests need more than a day. Hence, publishing software in Bioconda or updating already existing packages can be accomplished typically within minutes to a few hours.

Reproducible software management and distribution is enhanced by other current technologies. Conda integrates itself well with environment modules (http://modules.sourceforge.net/), a technology used nearly universally across HPC systems. An administrator can use Conda to easily define software stacks for multiple labs and project-specific configurations. Popularized by Docker, containers provide another way to publish an entire software stack, down to the operating system. They provide greater isolation and control over the environment a software is executed in, at the expense of some customizability. Conda complements container-based approaches. Where flexibility is needed, Conda packages can be used and combined directly. Where the uniformity of containers is required, Conda can be used to build images without having to reproduce the nuanced installation steps that would ordinarily be required to build and install a software within an image. In fact, for each Bioconda package, our build system automatically builds a minimal Docker image containing that package and its dependencies, which is subsequently uploaded and made available via the Biocontainers project (da Veiga Leprevost et al., 2017). As a consequence, every built Bioconda package is available not only for installation via Conda, but also as a container via Docker, Rkt (https://coreos.com/rkt), and Singularity (Kurtzer et al., 2017), such that the desired level of reproducibility can be chosen freely (Grüning et al., 2017).

## Discussion

By turning the arduous and error-prone process of installing bioinformatics software, previously repeated endlessly by scientists around the globe, into a concerted community effort, Bioconda frees significant resources to instead be invested into productive research. The new simplicity of deploying even complex software stacks with strictly controlled software versions enables software authors to safely rely on existing methods. Where previously the cost of depending on a third party tool - requiring its installation and maintaining compatibility with new versions - was often higher than the effort to re-implement its methods, authors can now simply specify the tool and version required, incurring only negligible costs even for large requirement sets.

For reproducible data science, it is crucial that software libraries and tools are provided via an easy to use, unified interface, such that they can be easily deployed and sustainably managed. With its ability to maintain isolated software environments, the integration into major workflow management systems and the fact that no administration privileges are needed, the Conda package manager is the ideal tool to ensure sustainable and reproducible software management. With Bioconda, we unlock Conda for the life sciences while coordinating closely with other related projects such as conda-forge and Biocontainers. Bioconda offers a comprehensive resource of thousands of software libraries and tools that is maintained by hundreds of international contributors. Although it is among the youngest, it outperforms all competing projects by far in the number of available packages. With almost six million downloads so far, Bioconda packages have been well received by the community. We invite everybody to participate in reaching the goal of a central, comprehensive, and language agnostic collection of easily installable software by maintaining existing or publishing new software in Bioconda.

## Acknowledgements

We thank the participants of various hackathons (e.g., the GalaxyP and IUC contribution fest, ELIXIR BioContainers and NETTAB hackathon) for porting numerous packages to Bioconda.

## Contributions

Kyle Beauchamp, Christian Brueffer, Brad Chapman, Ryan Dale, Florian Eggenhofer, Björn Grüning, Johannes Köster, Elmar Pruesse, Martin Raden, Jillian Rowe, Devon Ryan, Ilya Shlyakter, Andreas Sjödin, Christopher Tomkins-Tinch, and Renan Valieris (in alphabetical order) have written the manuscript. Johannes Köster and Ryan Dale have conducted the data analysis. Dan Ariel Sondergaard contributed by supervising student programmers on contributing recipes and maintaining the connection with ELIXIR. All other authors have contributed or maintained recipes.

## Online Methods

### Security Considerations

Using Bioconda as a service to obtain packages for local installation entails trusting that (a) the provided software itself is not harmful and (b) it has not been modified in a harmful way. Ensuring (a) is up to the user. In contrast, (b) is handled by our workflow. First, source code or binary files defined in recipes are checked for integrity via MD5 or SHA256 hash values. Second, all review and testing steps are enforced via the GitHub interface. This guarantees that all packages have been tested automatically and reviewed by a human being. Third, all changes to the repository of recipes are publicly tracked, and all build and test steps are transparently visible to the user. Finally, the automatic parts of the development workflow are implemented in the open-source software *bioconda-utils* (https://github.com/bioconda/bioconda-utils). In the future, we will further explore the possibility to sign packages cryptographically.

### Software management with Conda

Via the Conda package manager, installing software from Bioconda becomes very simple. In the following, we describe the basic functionality assuming that the user has access to a Linux or macOS terminal. After installing Conda, the first step is to set up the Bioconda channel via:

~~~
$ conda config --add channels conda-forge
$ conda config --add channels bioconda
~~~

Now, all Bioconda packages are visible to the Conda package manager. For example, the software CNV- kit (Talevich et al., 2016), can be searched for with

~~~
$ conda search cnvkit
~~~

in order to check if and in which versions it is available. It can be installed with:

~~~
$ conda install cnvkit
~~~

CNVkit needs various dependencies from Python and R, which would otherwise have to be installed in separate manual steps (Fig. 2a). Furthermore, Conda enables updating and removing all these dependencies via one unified interface. A key value of Conda is the ability to define isolated, shareable software environments. This can happen ad-hoc, or via YAML (https://yaml.org) files. For example, the following defines an environment consisting of Salmon (Patro et al., 2017) and DESeq2 (Love et al., 2014):

channels:

- bioconda
- conda-forge
- defaults

dependencies:

- bioconductor-deseq2 =1.16.1
- salmon =0.8.2
- - r-base =3.4.1

Given that the above environment specification is stored in the file env.yaml, an environment my-env meeting the specified requirements can be created via the command:

~~~
$ conda env create --name my-env --file env.yaml
~~~

To use the commands installed in this environment, it must first be “activated” by issuing the following command:

~~~
$ source activate my-env
~~~

Within the environment, R, Salmon, and DESeq2 are available in exactly the defined versions. For example, salmon can be executed with:

~~~
$ salmon --help
~~~

It is possible to modify an existing environment by using conda update, conda install and conda remove. For example, we could add a particular version of Kallisto (Bray et al., 2016) and update Salmon to the latest available version with:

~~~
$ conda install kallisto=0.43.1
$ conda update salmon
~~~

Finally, the environment can be deactivated again with:

~~~
$ source deactivate
~~~

### How isolated software environments enable reproducible research

With isolated software environments as shown above, it is possible to define an exact version for each package. This increases reproducibility by eliminating differences due to implementation changes. Note that above we also pin an R version, although the latest compatible one would also be automatically installed without mentioning it. To further increase reproducibility, this pattern can be extended to all dependencies of DESeq2 and Salmon and recursively down to basic system libraries like zlib and boost (https://www.boost.org). Environments are isolated from the rest of the system, while still allowing interaction with it: e.g., tools inside the environment are preferred over system tools, while system tools that are not available from within the environment can still be used. Conda also supports the automatic creation of environment definitions from already existing environments. This allows to rapidly explore the needed combination of packages before it is finalized into an environment definition. When used with workflow management systems like Galaxy (Afgan et al., 2016), bcbio-nextgen (https://github.com/chapmanb/bcbio-nextgen), and Snakemake (Köster and Rahmann, 2012) that interact directly with Conda, a data analysis can be shipped and deployed in a fully reproducible way, from description and automatic execution of every analysis step down to the description and automatic installation of any required software.

### Data analysis

The presented figures and numbers have been generated via a fully automated, reproducible Snakemake (Köster and Rahmann, 2012) workflow that is freely available under https://github.com/bioconda/ bioconda-paper.

